# The Effects of Negative Periocular Pressure on Biomechanics of the Optic Nerve Head and Cornea: A Computational Modeling Study

**DOI:** 10.1101/2022.05.17.492338

**Authors:** Babak N. Safa, Adam Bleeker, John P. Berdahl, C. Ross Ethier

**Affiliations:** Wallace H. Coulter Department of Biomedical Engineering, Georgia Institute of Technology/Emory University, Atlanta, GA, USA; Dean McGee Eye Institute Department of Ophthalmology, University of Oklahoma Health Sciences Center, Oklahoma City, OK, USA; Vance Thompson Vision, Sioux Falls, SD, USA

**Keywords:** Glaucoma, Biomechanics, IOP, Multi-pressure dial system, Finite Element Method

## Abstract

**Purpose:** To evaluate the effects of negative periocular pressure (NPP), and concomitant intraocular pressure (IOP) lowering, on the biomechanics of the optic nerve head (ONH) and cornea.

**Methods:** We developed a validated finite element (FE) model of the eye to compute tissue biomechanical strains induced in response to NPP delivered using the Multi-Pressure Dial (MPD) system. The model was informed by clinical measurements of IOP lowering and was based on published tissue properties. We also conducted sensitivity analyses by changing pressure loads and tissue properties.

**Results:** Application of -7.9 mmHg NPP decreased strain magnitudes in the ONH by c. 50% while increasing corneal strain magnitudes by c. 25%. Comparatively, a similar increase in corneal strain was predicted to occur due to an increase in IOP of 4 mmHg. Sensitivity studies indicated that NPP lowers strain in the ONH by reducing IOP and that these effects persisted over a range of tissue stiffnesses and spatial distributions of NPP.

**Conclusions:** NPP is predicted to considerably decrease ONH strain magnitudes. It also increases corneal strain but to an extent expected to be clinically insignificant. Thus, using NPP to lower IOP and hence decrease ONH mechanical strain is likely biomechanically beneficial for glaucoma patients.

**Translational Relevance:** This study provides the first description of how NPP affects ONH biomechanics and explains the underlying mechanism of ONH strain reduction. It complements current empirical knowledge about the MPD system and guides future studies of NPP as a treatment for glaucoma.

## Introduction

Glaucoma is the leading cause of irreversible blindness worldwide and is frequently associated with elevated intraocular pressure (IOP) [1]. The biomechanical response of certain ocular tissues is thought to play a significant role in glaucoma pathogenesis; specifically, elevated IOP causes supra-physiologic mechanical loading of the optic nerve head (ONH) tissues, i.e., lamina cribrosa (LC) and prelaminar tissue (PLT), which has been hypothesized to contribute to retinal ganglion cell (RGC) axonal damage and subsequent vision loss [2]– [4]. Clinical treatment for glaucoma seeks to decrease IOP [5]. Although current treatments can manage elevated IOP, vision loss continues to progress in almost 45% of patients, despite treatment [6], [7]. Further IOP lowering in patients with normal IOP who are progressing has been shown to slow progression, with most of these patients showing IOP acrophase at night [8], [9]. However, safely lowing IOP in these patients has proven difficult as the most effective treatments also carry significant morbidity [10], emphasizing the need for novel treatment options.

The Multi-Pressure Dial system (MPD; Equinox Ophthalmic, Inc., Newport Beach, CA) is a medical device that lowers IOP non-invasively via application of negative periocular pressure (NPP) [11], [12]. The MPD system consists of a pair of goggles connected to a programmable vacuum pump delivering adjustable NPP. Although experimentally undetected thus far, the negative pressure very likely increases globe volume. Consequently, IOP decreases as measured in experiments on a cadaveric model [13] and in living subjects [14], as well as predicted via a lumped-parameter biomechanical model [15]. Of note, we here define IOP as the pressure inside the eye referenced to the atmosphere (not to goggle pressure), consistent with all other pressure measurements in the body.

Despite IOP lowering due to NPP, there is no information regarding the effects of NPP on ocular tissue biomechanics, notably the ONH and cornea. Thus, our objective was to evaluate the biomechanical effects of NPP using a validated finite element (FE) modeling, a computational technique that allows efficient and flexible parametric investigation of the effects of mechanical loads on ocular tissues [16]–[22]. We focused on the ONH since it is an early and important site of neurological damage in glaucoma, and decreased ONH strain is likely beneficial for the treatment of glaucoma while increased strain of the ONH could be detrimental[3], [23], [24] [3], [23], [24]. Secondarily, we explored mechanical strains in the cornea induced by NPP, which is directly loaded by the application of NPP.

## Methods

We quantified the effects of NPP by simulating the biomechanical behavior of a human eye in several situations. We provide an overview of the simulations here, followed by full details below.

1. *Normotensive case*: A normal eye with an IOP of 15.8 mmHg.
2. *Goggle case*: The eye in Case 1, acted on by NPP distributed over the cornea and limbal region, decreasing gradually towards the posterior pole. IOP was reduced concomitantly, as is experimentally observed to occur during goggle wear [14].
3. *Hypertensive case:* The eye in Case 1, except that IOP was doubled to 31.6 mmHg.
4. *IOP fixed case:* Similar to Case 2, except that IOP was held constant at the normotensive level. Although it is known that IOP is lowered by NPP, this case is useful for understanding the mechanism by which NPP affects ONH biomechanics.

The biomechanical properties (stiffnesses) of ocular tissues were based on literature values, with adjustment to ensure that the modeled eye matched population-averaged ocular compliance data. We used ocular compliance for validation since it is a relatively well-characterized descriptor of corneoscleral shell biomechanical behavior, which in turn is expected to strongly influences how NPP will affect tissue strains within ocular tissues.

Tissue mechanical properties vary from one person to the next, so the biomechanical response due to NPP will vary from one person to the next. We, therefore, conducted a simplified sensitivity analysis to investigate the effects of such physiological variability. More complete sensitivity analyses are possible (Feola et al., 2016; Schwaner et al., 2020), but were beyond the scope of this initial study.

### Ocular geometry and FE modeling technical details

Our finite element model was based on existing FE models of the human eye [17], [22], with additional model dimensions obtained from the literature (Table 1). Following previous approaches [25], [26], we assumed axisymmetry to allow more rapid computations, considering a 5º “wedge” rather than an entire globe (Figure 1). We modeled the rectus muscles as a rigid body at the superior aspect of the globe, obliquely attached to the sclera (Figure 1), with the insertion site placed c. 6 mm posterior to the limbus, spanning approximately 1 mm [27].

**Table 1:** Summary of key dimensions for the eye model.

|  | Value (mm) | Source |
| --- | --- | --- |
| Corneal radius | 7.80 | [26], [48] |
| Corneal thickness | 0.56 | [26], [49] |
| Scleral radius | 11.00 | [22], [48] |
| Scleral thickness | 0.80 | [17], [22] |
| Retinal thickness | 0.20 | [17], [22] |
| LC Thickness | 0.20 | (Feola et al., 2016; Sigal et al., 2005) |

**Figure 1:**
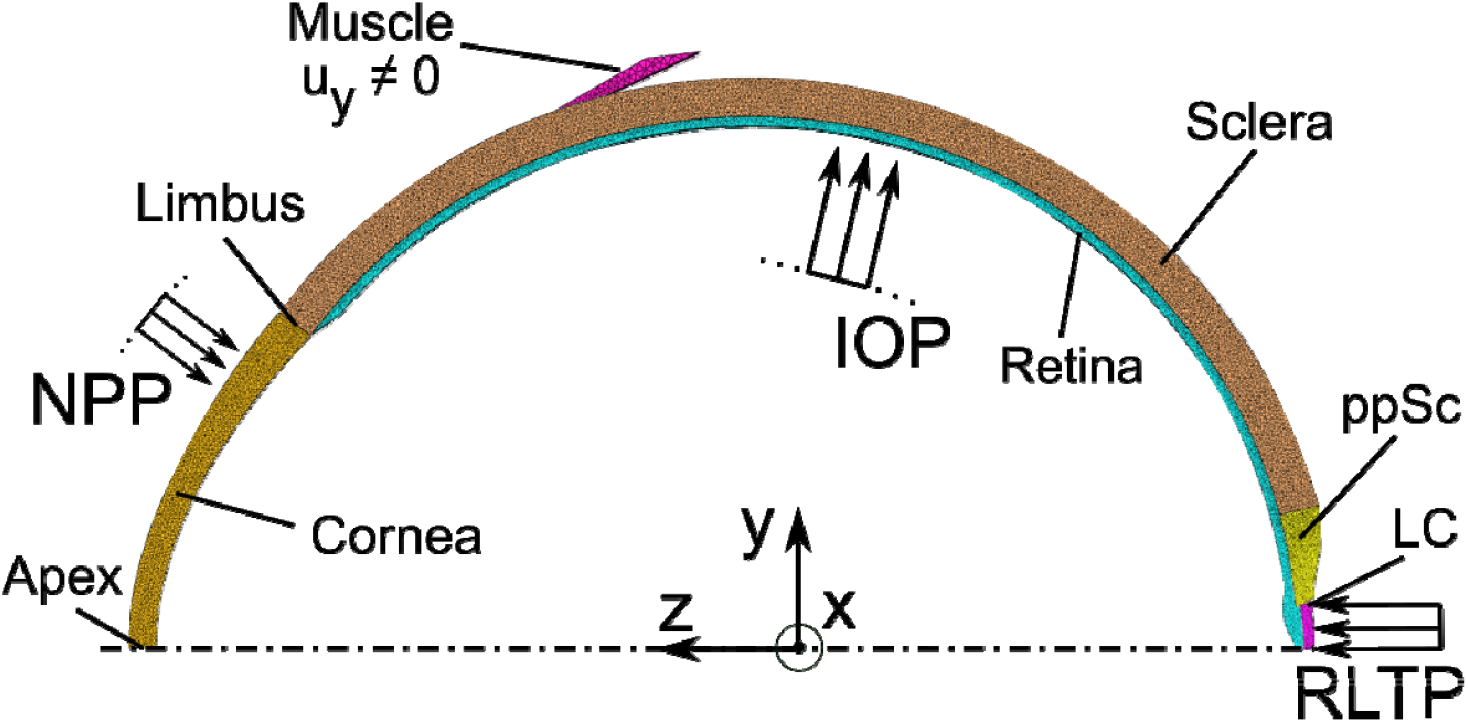
The finite element model used in this study, including the corneoscleral shell and ONH (ppSC=peripapillary sclera; LC=lamina cribrosa). The extraocular rectus muscle attachment is also shown, which is modeled as a rigid body free to move along the y-axis. The figure also schematically represents the loads, namely intraocular pressure (IOP), negative periocular pressure (NPP), and retrolaminar tissue pressure (RTLP).

This geometry was meshed with conforming second-order tetrahedral elements (TET10) in Gmsh (v4.8.4) [28] using the “Frontal” algorithm [29]. Following initial mesh generation, 10 smoothing steps and 10 mesh optimization steps were performed, and the mesh was then exported as a Version 2 ‘msh’ file (ASCII). Based on a preliminary mesh sensitivity analysis, we used 446,528 elements (element size factor c. 25-100 μm) to tesselate the domain, with elemental density enhanced 4-fold in the ONH.

All FE simulations were carried out using FEBio v3.5.1 (Maas et al., 2012). We enforced axisymmetry through the appropriate specification of nodal degrees of freedom. Specifically, nodes on the axis of symmetry lying within the cornea, retina, and LC were constrained to move along the z-axis, while nodes on the bounding planes of the wedge were constrained to move in-plane, so their displacement was radial relative to the axis of symmetry (Figure 1). The muscle was free to move along the y-axis (*u*_*y*_ ≠ 0), but was constrained in the remaining five degrees of freedom (Figure 1). To check the impact of this boundary condition, we mimicked a hinge-like muscle connection by allowing the muscle to also rotate around the x-axis (see supplementary Figure S1); however, this modification did not result in any significant change in the mechanical response of the model, and thus for simplicity, we returned to the original boundary condition (only y-displacements allowed).

### Specification of ocular pressures

#### Normotensive case

This case describes a normal eye without goggles with the following loads typical for a healthy eye: IOP = 15.8 mmHg, NPP = 0, and retrolaminar pressure (RLTP) = 8 mmHg [13]. IOP was uniformly applied on the interior surface of the corneoscleral shell, while the NPP was applied uniformly from the corneal apex to ∼2.5 mm posterior to the limbus, then decreasing linearly to zero near the edge of the ppSC. Unfortunately, there are no experimental data regarding the spatial distribution of NPP; therefore, to assess the effect of uncertainty in the spatial distribution of NPP, we also varied the distribution of NPP on the globe surface (see Supplementary Figure S4), which showed no significant difference in results.

#### Goggle case

To test the effects of NPP, we modeled a normotensive eye with MPD goggles applied, specifying the following pressures: IOP = 11.5 mmHg, NPP = -7.9 mmHg, and RLTP = 8 mmHg. NPP and IOP were based on experimental measurements [14], where it was observed that imposing an NPP equal to 50% of the baseline IOP (here, NPP = -7.9 mmHg) reduced IOP to 11.5 mmHg.

#### Hypertensive Case

We simulated the effects of elevated IOP (31.6 mmHg), with all other inputs being the same as in the Normotensive case. This allowed us to compare strains observed in the Goggle case to those occurring in ocular hypertension.

#### IOP fixed case

NPP both lowers IOP and expands the corneoscleral shell, which could be expected to have opposite effects on strains in the ONH [25]. Thus, to investigate the effects of only NPP-induced corneoscleral shell expansion on ONH biomechanics, we held IOP fixed at 15.8 mmHg while imposing NPP. This would likely not occur clinically but provides a useful mechanistic understanding of the effects of NPP on the eye’s biomechanics.

### Specification of tissue biomechanical properties

To investigate the effects of tissue stiffness, we considered a range of tissue mechanical properties, which we refer to as *tissue parameter sets*. In all cases except those noted explicitly below, we used a nearly-incompressible hyperelastic solid constitutive formulation to describe the mechanical behavior of tissues, where Young’s modulus *E* was the model parameter (for details of constitutive equation see Appendix).

In the *Baseline tissue parameter set*, values of *E* were based on literature reports, with modest adjustments to match clinical ocular compliance data (Table 2). To account for the microarchitecture of the peripapillary sclera (ppSC), we included circumferential (along the x-axis; Figure 1) collagen fibers in the ppSC, described by a 1-D linear elastic constitutive relation, with fiber modulus value (*E*_*f*_ = 41.83 MPa) (Grytz et al., 2014). When modeling fibers, we did not consider the effect of fiber distribution and crimp, which effectively corresponds to an upper bound on ppSC stiffness. Lastly, as a diagnostic test case, we considered the effects of reducing ppSC stiffness by eliminating collagen fibers from the ppSC, similar to Sigal et al. (Sigal et al., 2005, 2004). This corresponds to a lower bound for ppSC stiffness and produced results in agreement with the previous work by Sigal et al. (Figure S3).

**Table 2:** Summary of tissue biomechanical stiffnesses (*E*, in MPa) used in simulations, with corresponding [literature reference]. In all simulations, the muscle was treated as rigid. “Same” indicates that the values are the same as those listed for the *Baseline tissue parameter set*.

| Tissue parameter sets |  |  |  |
| --- | --- | --- | --- |
| Tissue | Baseline | Soft shell/Stiff shell | Soft LC/Stiff LC |
| Sclera | 4.50 [1.5 □ (Sigal et al., 2005)] | 2.25/9 | Same |
| ppSC matrix | 4.50 [1.5 □ (Sigal et al., 2005)] | 2.25/9 | Same |
| ppSC fibers | 41.83 [(Grytz et al., 2014)] | 20.92/83.66 | Same |
| LC | 0.30 [(Sigal et al., 2005)] | Same | 0.0026/3 |
| Retina | 0.06 [(Feola et al., 2016)] | Same | Same |
| Cornea | 0.84 [(Blackburn et al., 2019)] | 0.42/1.68 | Same |

#### Soft and Stiff (corneoscleral) shell tissue parameter sets

We investigated the effects of corneoscleral shell stiffness by doubling or halving sclera, ppSC, and cornea matrix stiffnesses, denoted as *Stiff shell* and *Soft shell tissue parameter sets*, respectively (Table 2). This range of variation was inspired by the reported range of values for the human eye’s mechanical properties [22], [30]–[32].

#### Soft and Stiff LC tissue parameter sets

We also studied the effect of varying the stiffness of the LC. In the *Stiff LC tissue parameter set*, we set the stiffness of the LC to be similar to scleral stiffness (*E* = 3 Mpa), which represents a (non-physiological) limiting high-stiffness case, useful for sensitivity analysis (Table 2). The *Soft LC parameter set* was informed by *ex vivo* biomechanical measurements of the LC (Safa et al., 2021). Specifically, we modeled the LC as a compressible neo-Hookean material with *E* = 2.6 kpa and Poisson’s ratio *v* = 0.23 following a well-established formulation [33].

### Model validation

To validate the FE model, we compared our model’s ocular compliance to literature values. Specifically, we computed the compliance of the corneoscleral shell in the Normotensive case as *ϕ* = d(IOV)/d(IOP), where d(IOV) is the change in intraocular volume, i.e., the volume enclosed within the corneoscleral shell (Figure 1), in response to a change in IOP, *d*(*IOP*). The computed value of *ϕ* was compared to previous values summarized from multiple *in vivo* experiments [34] (Table 3), using the formula of McEwen and Helen: *ϕ* = (*a* × IOP + *b*)^−1^, where *a =* 0.015 – 0.027 μL^−1^, and *b =* 0.03 – 0.31 mmHg/ μL evaluat ed at an IOP of 15 mmHg. To match compliance values in the literature we increased the stiffness of the sclera, cornea, and ppSC matrix used by Sigal et al. [22] by a factor of 1.5 (Table 2).

### Modeling outcomes and data analysis

Our primary outcome measures were the nodal distribution of the first and third principal Lagrangian strain (*E*_I_ and *E*_III_, respectively), two standard measures of tissue deformation, representing maximal tensile (*E*_I_) and compressive (*E*_III_) strains in an isochoric deformation. Note that due to the compressibility of the LC in the *Soft LC parameter set*, negative *E*_*I*_ values occurred, which were visualized using a value-preserving bi-symmetric logarithmic transformation (Perrotta, 2022; Webber, 2013).

We considered strains in four tissue regions of interest: the LC (1643 LC nodes on the *x* = 0 plane), the prelaminar tissue (2357 PLT nodes on the *x* = 0 planes), the limbus (240 nodes shared by the cornea and sclera in 3D), and the corneal apex (440 corneal nodes on the *x* = 0 plane) (Figure 1). More specifically, the PLT was defined as tissue within the retina having a radial distance from the axis of symmetry less than the LC radius in the *x-*y plane. The corneal apex was defined as the corneal region centered on the symmetry plane having a diameter of 3.06 mm, which matches the region that would be directly applanated during Goldmann tonometry [35], a region that is clinically familiar to practitioners. Finally, we evaluated the uniformity of the spatial placement of the selected nodes to avoid bias due to potential nodal clustering, confirming the appropriateness of our sampling (see supplementary Figure S2).

## Results

### Model validation

We expected the eye’s biomechanical response to NPP to depend strongly on volume increase in the corneoscleral shell, which can be macroscopically characterized by ocular compliance, *ϕ*. It was, therefore, important to validate our model against the values of *ϕ* provided in the literature. The compliance of our model in the Normotensive case was *ϕ* = 3.1μL/mmHg, which is within the range of reported values for the human eye from *in vivo* experiments (*ϕ* = 1.4−3.9μL/mmHg [34]).

### Effects of NPP

#### Normotensive case vs. Goggle case

It was of interest to probe the strains in the LC, PLT, limbus, and at the corneal apex due to NPP in a normotensive eye (i.e., comparison between the Normotensive and Goggle cases). Using the *Baseline tissue parameter set*, NPP caused the median *E*_I_, a measure of tissue tension, to decrease by 53.9% in the LC, from 0.52% [0.40%, 0.73%] (median [interquartile range]; Figure 2 A and M) to 0.24% [0.18%, 0.34%] (Figure 2 D and M). Similarly, in the PLT, *E*_I_ decreased by 55.3%, from 0.92% [0.78%, 1.13%] (Figure 2 A and M) to 0.41% [0.35%, 0.50%] (Figure 2 D and M). Conversely, *E*_I_ at the limbus increased by 23.7%, from 0.69% [0.67%, 0.71%] (Figure 2 B and M) to 0.86% [0.83%, 0.88%] (Figure 2 E and M), and at the corneal apex by 25.3%, from 0.95% [0.74%, 1.20%] (Figure 2 C and M) to 1.19% [0.94%, 1.48%] (Figure 2 F and M).

**Figure 2:**
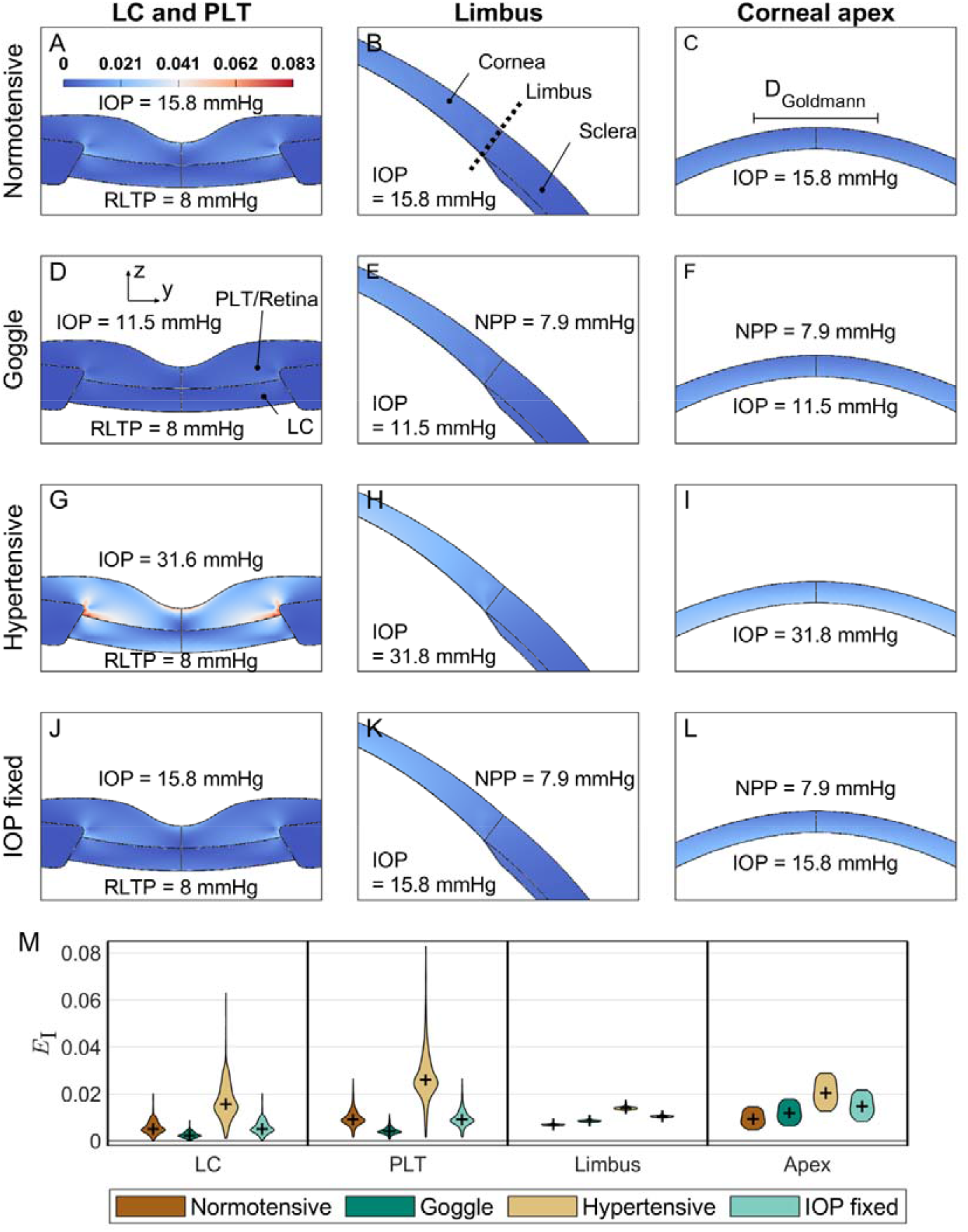
(**A-L**) Maps of the first principal Lagrangian strain (), a measure of tissue stretching, in the LC, PLT, limbus, and at the corneal apex for tissue stiffnesses defined by the *Baseline tissue parameter set*. Each of the top four rows correspond to one case (i.e., Normotensive [**A-C**], Goggle [**D-F**], Hypertensive [**G-I**], and IOP fixed [**J-L**]). The bottom row (**M**) provides a summary of values (violin plots, with median shown by ‘+’) in each region. D_Goldmann_ = 3.06 mm is the diameter of the applanated region during Goldmann tonometry, which is a clinically familiar region that we used to define the corneal apex region.

Similarly, NPP caused the magnitude of *E*_III,_, a measure of tissue compression, to decrease in the LC by 52.8%, with the value of *E*_III_ changing from -0.79% [-1.06%, -0.56%] (Figure 3 A and M) to -0.37% [-0.50%, -0.25%] (Figure 3 D and M), while in the PLT the magnitude of *E*_III_ decreased by 54.4%, with the value of *E*_III_ changing from -0.68% [-0.93%, -0.52] (Figure 3 A and M) to -0.31% [-0.43%, -0.24] (Figure 3 D and M). Contrary to the LC and PLT, NPP caused the magnitude of *E*_III_ at the limbus to increase by 24.2%, with *E*_III_ changing from -0.86% [-0.98%, -0.79%] (Figure 3 B and M) to -1.07% [-1.21%, -0.97%] (Figure 3 E and M), and at the corneal apex by 24.8%, with *E*_III_ changing from -1.78% [-2.20%, -1.41%] (Figure 3 C and M) to -2.22% [-2.71%, -1.78%] (Figure 3 F and M).

**Figure 3:**
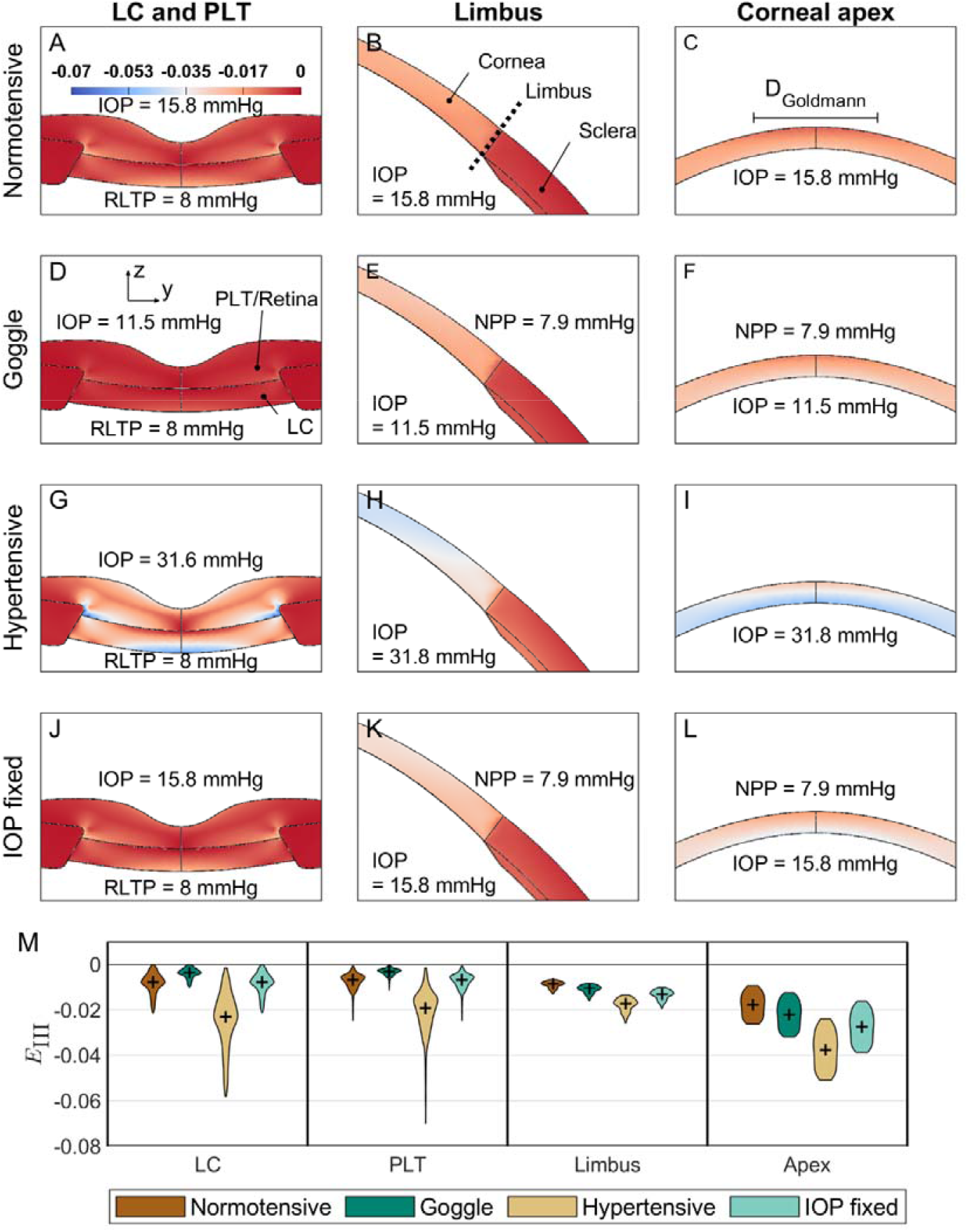
(**A-L**) Maps of the third principal Lagrangian strain (), and (**M**) summary of values for tissue stiffnesses defined by the *Baseline tissue parameter set*. For a detailed description of each panel see the caption of Figure 2.

#### Hypertensive case

In the Hypertensive case, where IOP was set to 31.6 mmHg and no goggles were present, the median *E*_I_ was significantly higher in the LC and PLT relative to the Normotensive case (201.2% and 184.0% greater in the LC and PLT, respectively, using the *Baseline tissue parameter set*; Figure 2 G and M). Similarly, median *E*_I_ was 101.4% and 115.9% greater in the limbus and at the corneal apex, respectively (compared to the Normotensive case; Figure 2 H, I, and M). We note that the strain increases in the cornea due to ocular hypertension were almost four times larger than the 25% strain increases observed in the Goggle case (all referenced to the Normotensive case). Similarly, the magnitude of *E*_III_ increased by 190.9% and 180.1% in the LC and PLT, respectively (compared to the Normotensive case; Figure 3 G and M). In addition, the magnitude of *E*_III_ increased by 101.0% at the limbus, and by 111.3% at the corneal apex (Figure 3 H, I, and M, using the *Baseline tissue parameter set*), which were again approximately four times larger than the 25% corneal strain increases observed in the Goggle case.

#### IOP fixed case

In this case, the IOP was artificially held constant when NPP was applied. We observed that NPP had minimal effects on *E*_I_ and *E*_III_ magnitudes in the LC and PLT, with a difference of less than 1% compared to the Normotensive case (Figures 2 and 3, J-L and M). However, fixing the IOP led to a large increase in *E*_I_ in the cornea; specifically, at the limbus, *E*_I_ increased by 51.2% (Figure 2 K), and at the corneal apex it increased by 56.1% (Figure 2 I), all referenced to the Normotensive case. Similarly, the magnitude of *E*_III_ at the limbus increased by 51.5% (Figure 3 K), and by 54.5% at the corneal apex (Figure 3 I). In every instance, these corneal strain increases relative to the Normotensive case were much larger than those observed in the Goggle case.

### Sensitivity of tissue strains to stiffness modulation

#### Soft and Stiff shell tissue parameter set

Changing corneoscleral shell stiffness within a physiological range using the *Soft/Stiff Shell tissue parameter sets* (Table 2) had only a small effect on strain magnitudes in the LC and PLT. More specifically, relative to the Normotensive and Goggle cases that were modeled using the *Baseline tissue parameter set* of tissue properties, softening the shell caused an overall increase of 3.4-16.0% in LC and PLT strain magnitudes, while stiffening the corneoscleral shell caused less than 5% change in strain magnitudes (Figure 4 A, B, E, and F). In contrast, the strains at the limbus and corneal apex approximately doubled using the *Soft shell tissue parameter set* and halved for the *Stiff shell tissue parameter set* compared to results obtained with the *Baseline tissue parameter set* for both the Normotensive and Goggle cases (Figures 4 C, D, G, and H). Importantly, the decrease in ONH strains and increase in corneal strains seen in the Goggle case relative to the Normotensive persisted when using the *Soft/Stiff tissue parameter sets* to describe tissue stiffnesses (Figure 4).

**Figure 4:**
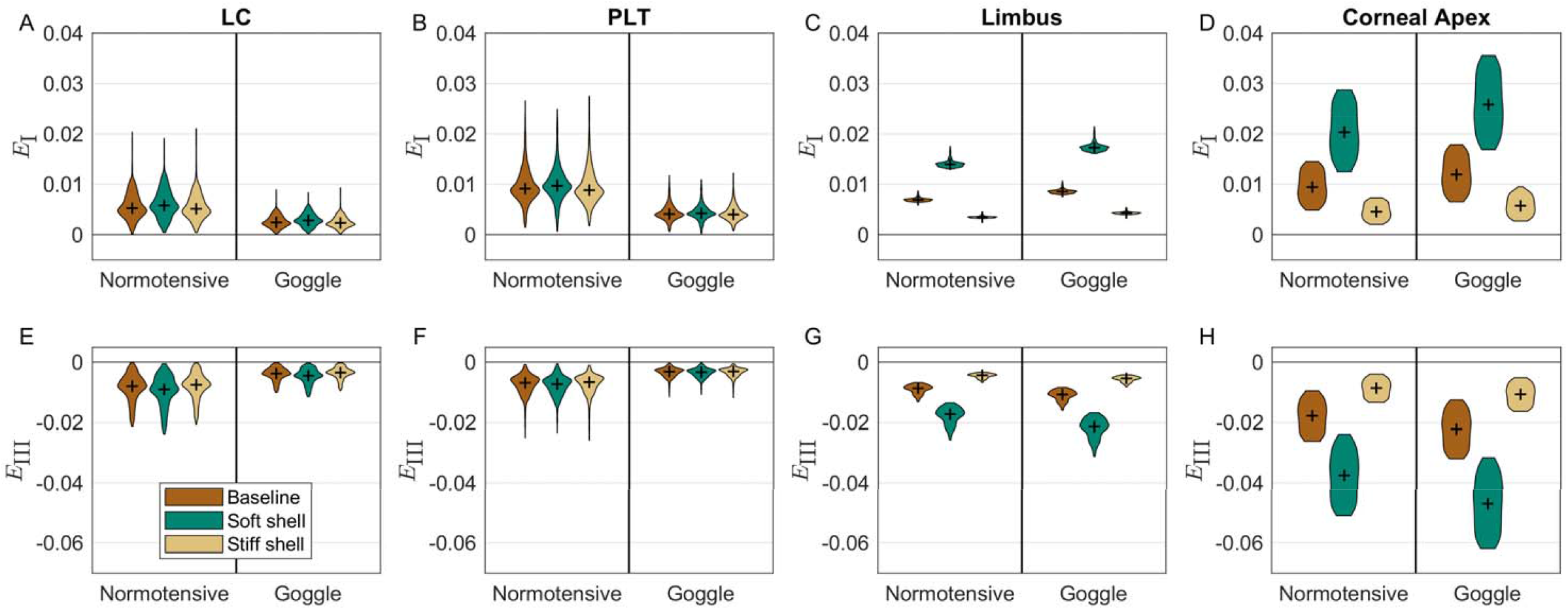
Changes in first (*E*_I_) and third (*E*_III_) principal strains due to varying corneoscleral shell stiffness. Calculations were carried out for both the Normotensive and Goggle cases, using the *Baseline, Soft Shell* and *Stiff Shell tissue parameter sets* (Table 2) Each panel depicts the strain values in the Normotensive and Goggle cases in different tissues (lamina cribrosa [LC; **A** and **E**], prelaminar tissue [PLT; **B** and **F**], limbus [**C** and **G**], and corneal apex [**D** and **H**]). Changing corneoscleral shell stiffness did not change the effect of NPP in decreasing strain magnitudes in the lamina cribrosa (LC; **A** and **E**) and prelaminar tissue (PLT; **B** and **F**); however, softening (stiffening) corneoscleral shell stiffness increased (decreased) *E*_I_ and *E*_III_ magnitudes at the limbus (**C** and **H**) and corneal apex (**D** and **H**). Data is shown using violin plots, with the median shown by ‘+’.

#### Soft and Stiff LC tissue parameter sets

As expected, significantly lowering the LC stiffness, and adding compressibility to mimic the *ex vivo* properties of the ONH (*Soft LC parameter set*; Table 2) markedly increased strain magnitudes in the LC and PLT (Figure 5 A and C). Specifically, in the Normotensive case simulated using the *Soft LC parameter set, E*_I_ of the LC increased by a factor of 6.1 relative to the Normotensive case simulated with the *Baseline parameter set*, where *E*_I_ increased by a factor of 2.8 in the PLT as a result of using *Soft LC parameter set* (Figure 5 A). Similarly, in the Goggle case, when using the *Soft LC parameter set, E*_I_ was greater by a factor of 5.7 in the LC and 2.9 in the PLT as compared to the *Baseline tissue parameter set* (Figure 5 C). Further, the magnitude of *E*_III_ was greater by a factor of 31.9 in the LC and 3.6 in the PLT in the Normotensive case, and by a factor of 64.9 in LC and 3.6 in the PLT in the Goggle case (Figure 5 A and C), again compared to using the *Baseline tissue parameter set*. Conversely, stiffening the LC (using the *Stiff LC tissue parameter set*; Table 2), so that the LC stiffness was similar to that of the sclera, decreased *E*_I_ and the magnitude of *E*_III_ in both the LC and PLT by c. 75% (Figure 5).

**Figure 5:**
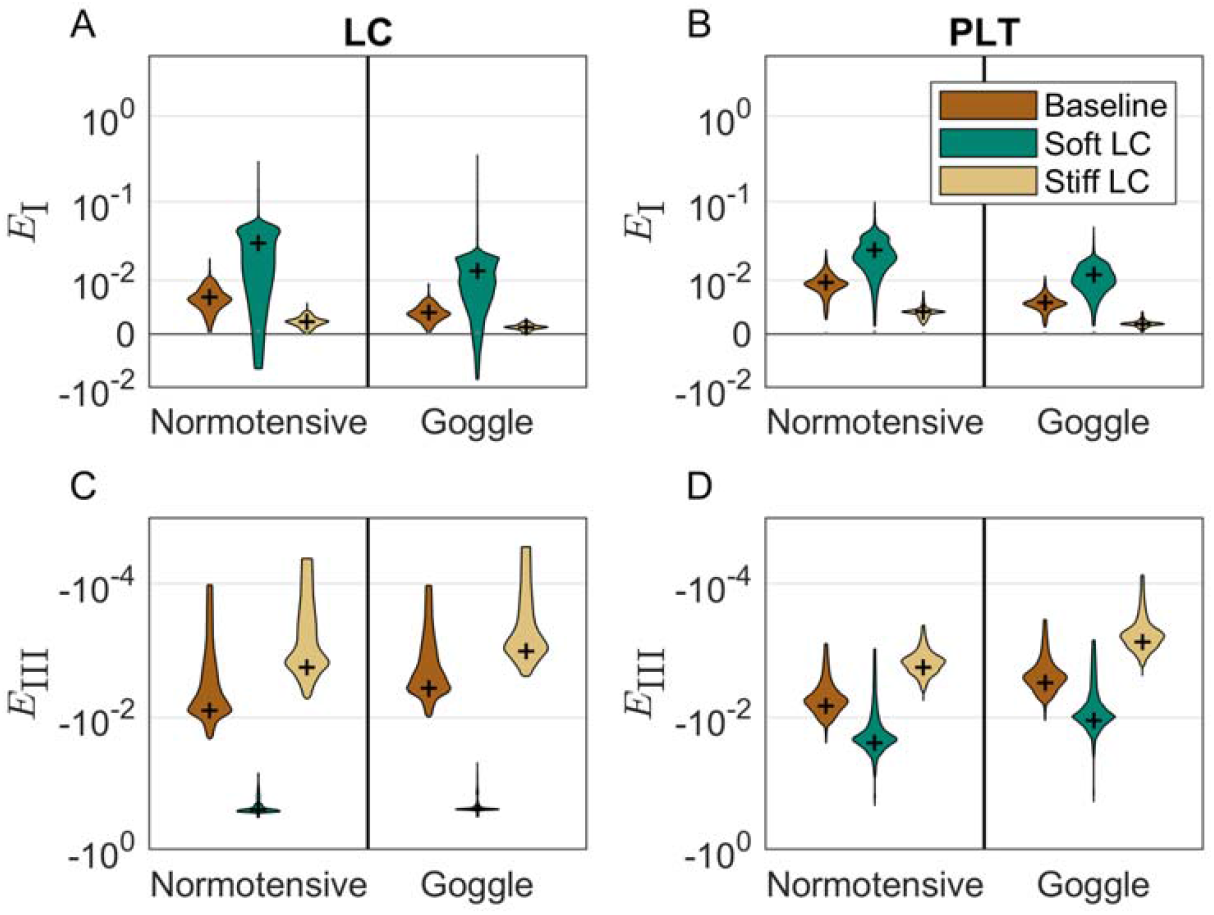
The effects of changing LC stiffness on strains (*E_I_* and *E_III_*) in the lamina cribrosa [LC; **A** and **C**] and prelaminar tissue [PLT; **B** and **D**]) using *Baseline, Soft LC, and Stiff LC tissue parameter sets* (Table 2). The magnitudes of *E_I_* and *E_III_* increased in both the LC (**A** and **B**) and the PLT (**C** and **D**) when using *Soft LC tissue material parameter set* relative to the values computed when using the *Baseline tissue material parameter set*. Conversely, for the Stiff LC case they decreased. However, despite these changes, for both the *Soft and Stiff LC material parameter sets*, the magnitudes of *E_I_* and *E_III_* decreased in the Goggle case relative to the Normotensive case. Data is shown using violin plots and the median is marked with ‘+’.

Interestingly, the observation that *E*_I_ and *E*_III_ magnitudes in the LC and PLT were lower in the Goggle case relative to the Normotensive case was largely unaffected by changing the stiffness and compressibility of the LC. The exception was for the values of *E*_III_ in the LC for the *Soft LC tissue parameter set*, where only a 3.9% decrease in LC *E*_III_ magnitude was observed due to Goggles compared to the Normotensive case (Figure 5 C).

## Discussion

This study evaluated the biomechanical response of the optic nerve head (ONH) and cornea to negative periocular pressure (NPP). Our results indicate that NPP significantly decreases mechanical strain in the ONH while increasing it in the cornea (Figures 2 and 3). This effect occurred over a wide range of tissue stiffnesses. Consistent with the known benefit of lowering IOP, and the understanding of the role of ONH biomechanics in glaucoma, reducing ONH biomechanical strain is predicted to slow or stop retinal ganglion cell axonal damage. This reduction is ONH strain is thus predicted to be beneficial for glaucoma patients.

The decrease in the ONH strains was not sensitive to changing the corneoscleral shell stiffness (LC, Figure 4 A and E; PLT Figure, 4 B and F), nor was it sensitive to changing the spatial distribution of NPP (Figure S4), suggesting that the decrease in IOP due to NPP is the primary cause of the decrease in the ONH strains (Figures 2 and 3). This is also consistent with the outcome of the IOP Fixed case, in which globe expansion without IOP lowering did not reduce ONH strains. This result was non-obvious since NPP both lowers IOP (expected to reduce ONH strain) and expands the corneoscleral shell (expected to increase ONH strain); it appears that for the range of tissue stiffnesses that we considered, the IOP decrease effect dominates any expansion of the corneoscleral shell.

The insensitivity of computed strains in the LC and PLT to a change in corneoscleral shell stiffness was due to including ppSC fibers in our model since these are an important determinant of ppSC deformations and hence scleral canal expansion during globe volume changes. In fact, when collagen fiber-reinforcement in the ppSC was eliminated, ONH strains were sensitive to changing corneoscleral shell stiffness (Figure S3), consistent with the findings of Sigal and co-workers, in which ppSC fibers were not considered [25]. Nevertheless, independent of whether ppSC collagen fibers were modeled or not, NPP decreased ONH strains, further supporting the strain-reducing effect of NPP on the ONH.

Interestingly, the biomechanical properties of the LC, and especially its compressibility, showed a significant effect on the strain-reducing effect of NPP in ONH (Figure 5). More specifically, tissue compressibility in the LC, observed in *ex vivo* [36] and *in situ* [37] experiments, significantly affected the ONH strains. For instance, using the *Soft LC tissue parameter set* significantly diminished the decrease in strain magnitudes from Reference to Goggle configurations (Figure 5). Therefore, it is likely that the patients with different ONH biomechanics and glaucoma types would have some variability in the magnitude of ONH strain reduction for a given decrease in IOP due to NPP. For example, in pseudoexfoliation glaucoma (PEXG), LC stiffness is decreased (Braunsmann et al., 2012), whereas in primary open-angle glaucoma (POAG), the LC tends to stiffen [2], [39], [40]. Despite this variability, reducing IOP with NPP is still likely to be beneficial since ONH strains were decreased even at extreme LC stiffness and softness.

Although NPP decreased strain magnitudes in the ONH, it nonuniformly increased them in the cornea (<1% strain; Figures 2 and 3). However, the increase in corneal strain induced by NPP was almost four-fold smaller than the increase in corneal strain observed when IOP was increased from 15.6 mmHg to 31.6 mmHg (in the absence of goggles; Hypertensive case; Figures 2 and 3). Based on these findings, the observed increase in corneal strain due to NPP is estimated to be equivalent to that due to an IOP increase of c. 4 mmHg in a normotensive eye. Although the role of corneal biomechanics in ocular pathophysiology remains an active area of vision science research, clinical experience indicates that the cornea tolerates high IOPs (c. 30 mmHg) relatively well, suggesting that strains equivalent to those due to an IOP increase of 4 mmHg would not cause significant sequelae in the cornea or angle. For example, Hjortdal demonstrated regional differences in meridional and circumferential strains in the human cornea produced by IOPs up to 100 mmHg [41]. Despite this significant amount of pressure loading and increase in the transcorneal pressure difference, no sign of damage was observed, which in collagenous soft tissues presents as a progressive decrease in tensile modulus [42], [43].

Further, several prospective clinical trials have examined the safety of NPP applied by the MPD system. A phase I trial exposed 30 eyes to an NPP of -15 mmHg for 30 minutes. No adverse events, including corneal events, were noted immediately following goggle removal and at 7 days post-intervention [44]. Similar safety outcomes were observed in a cohort of 65 healthy eyes treated with different amounts of NPP [14]. Additionally, no corneal adverse events were observed in glaucomatous eyes exposed to NPP overnight [11], [45]. Clinical evidence corroborates existing corneal biomechanics research suggesting that the corneal strain induced by NPP is likely clinically insignificant.

A potential limitation of this study was the assumption that the IOP decrease due to NPP was unaffected by changing corneoscleral stiffness (Figure 4) and the spatial distribution of NPP (Figure S4). In reality, we expect that corneoscleral stiffness and NPP distribution will modestly affect the magnitude of the IOP decrease; therefore, in future, it would be worthwhile extending the model to include blood flow and fluid-solid interactions. Further, for numerical efficiency, we assumed that the eye to be axisymmetric relative to its optical axis. In reality, the ONH axis is slightly off-center relative to the optical axis; however, this slight difference is unlikely to affect our results. Finally, we assumed the extraocular rectus muscle to be a rigid body, which is a simplification justified by the significantly higher stiffness of the rectus muscles and tendons relative to the other ocular tissues [46].

In conclusion, this study provides novel insights into the biomechanical effects of NPP on the ONH and cornea. We showed that NPP decreases strain in ONH tissues by reducing IOP, while NPP increases cornea strains. The decrease of ONH strains is likely beneficial for glaucoma patients, while NPP-induced corneal strains are relatively small and likely clinically insignificant.

## Acknowledgments

We acknowledge our funding sources: the Georgia Research Alliance (CRE) and the BrightFocus Foundation (postdoctoral fellowship G2021005F, BNS).

## Appendix Constitutive equations

Tissues were treated as incompressible neo-Hookean materials described by the following constitutive equation:

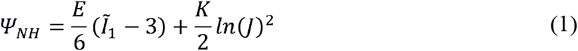

where *Ĩ*_1_ is the first invariant of the deviatoric right Cauchy-Green strain tensor 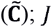 is the Jacobian of the deformation, *J* = det(**F**) with **F** being the deformation gradient tensor; *E* is the Young’s modulus; and *K* is the bulk modulus. The values of *E* were taken from the literature (Table 2). When specifying *K*, we note that the material incompressibility constraint (i.e., *J* = 1), nominally eliminates the effect of *K* on material behavior; however, *K* is used as a penalty factor to enforce incompressibility, and it was set as a large value relative to *E* (Weiss et al., 1996).

We implemented the effects of collagen fibers in the ppSC using an equation suitable for the uncoupled nearly incompressible formulation (modified from *fiber-pow-linear uncoupled* [47]):

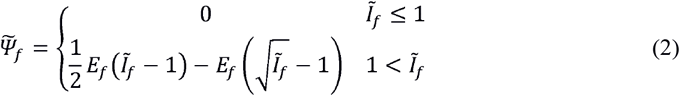

where *E*_*f*_ is the fiber Young’s modulus, and *Ĩ*_*f*_ is the square of fiber stretch, calculated as 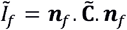, where ***n***_*f*_ is the unit vector defining the fiber orientation.

---

**Figure S1:**
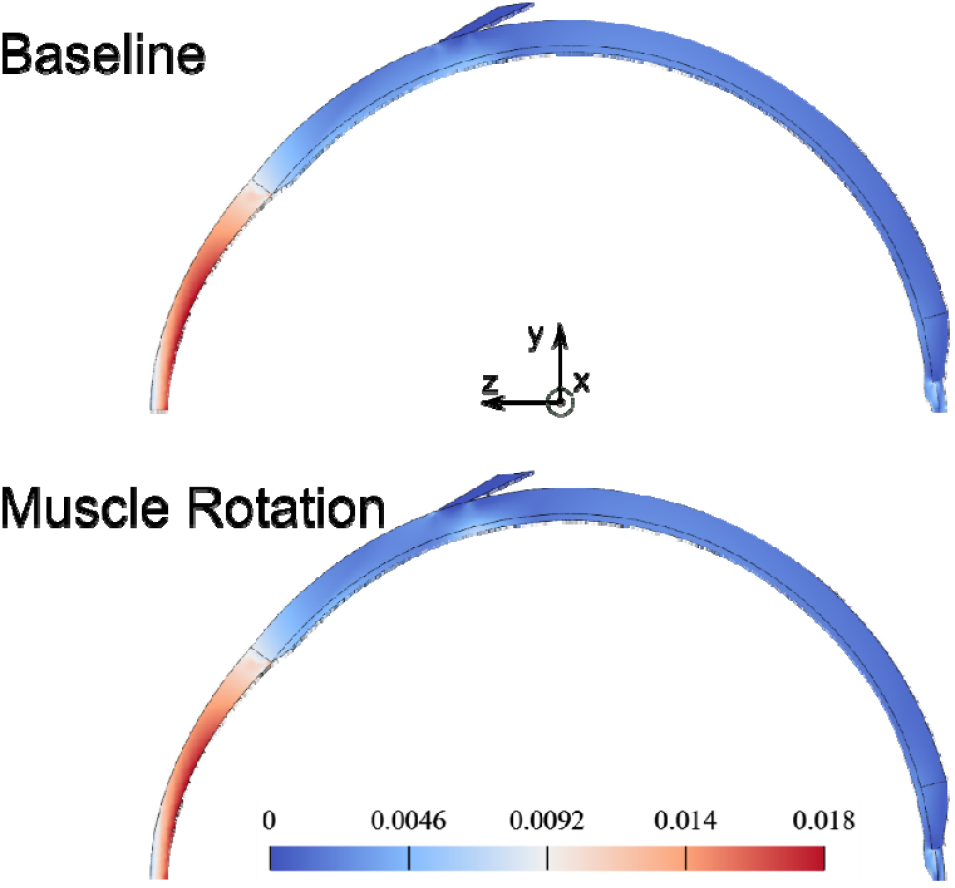
Plots of first principal Lagrangian strain,, in the Goggle case showing the effects of changing the muscle boundary condition. In the top row the muscle was only allowed to translate in the y-direction, while in the simulation shown in the bottom row, the muscle was also allowed to rotate around the x-axis. It is evident that the two cases showed almost identical strain distributions.

**Figure S2:**
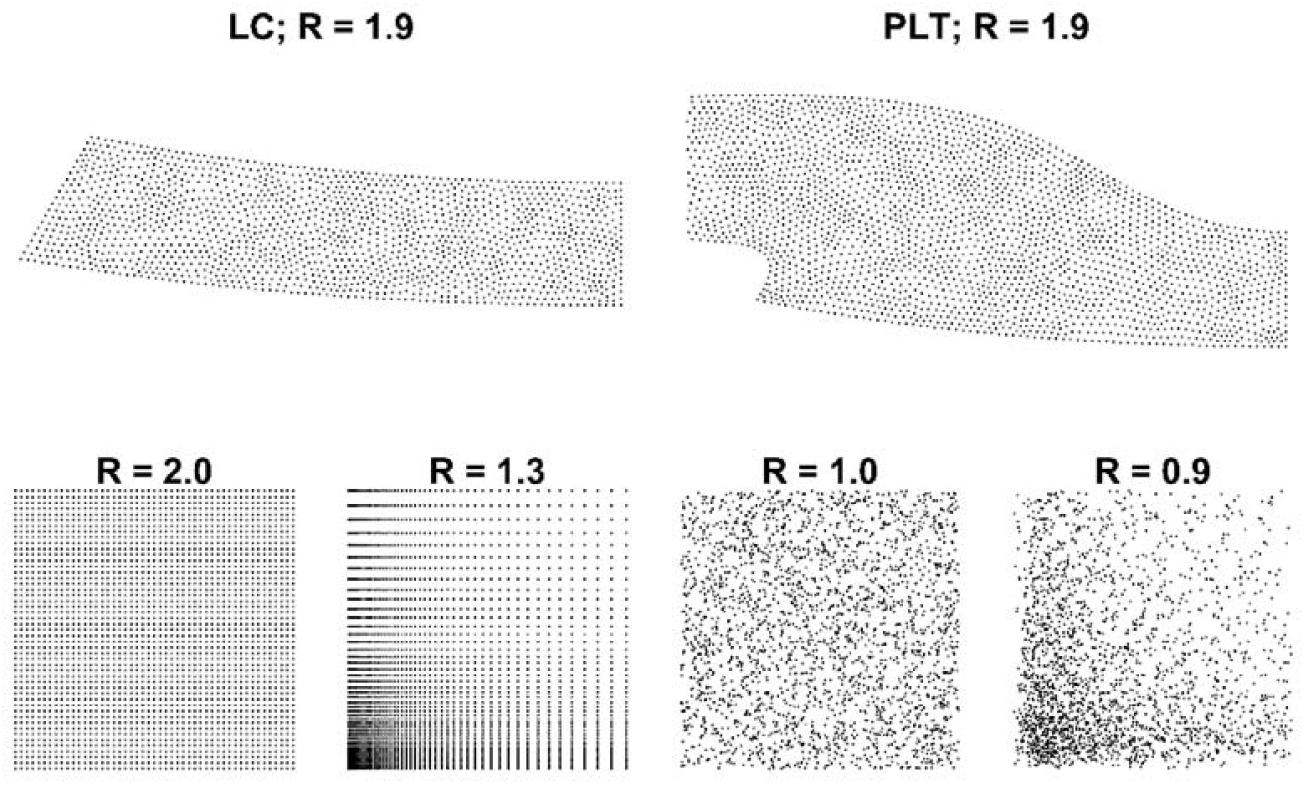
Quantification of the spatial uniformity of finite element nodes in the lamina cribrosa (LC) and prelaminar tissue (PLT), important since we report strain values at nodes and wish to avoid biasing our reported values by over-sampling a specific tissue sub-region. For this purpose, we use a quantitative measure commonly used in geoscience [50], defined as, where is the average distance from the nearest neighbor and is the expected statistical distance from the nearest point, P. The maximum theoretical value of *R* is 2.1491 for an infinitely large population of points having a hexagonal distribution (Clark and Evans, 1954). Here we calculate for the LC and PLT (first row), and for four other distributions with a similar number of nodes but different levels of nonuniformity, ranging from a uniform square grid (second row left) to a biased random distribution (second row right). It is evident that the values for the LC and PLT nodes are similar to the value for the uniform square grid. Thus, reporting strains at nodal location was deemed appropriate.

**Figure S3:**
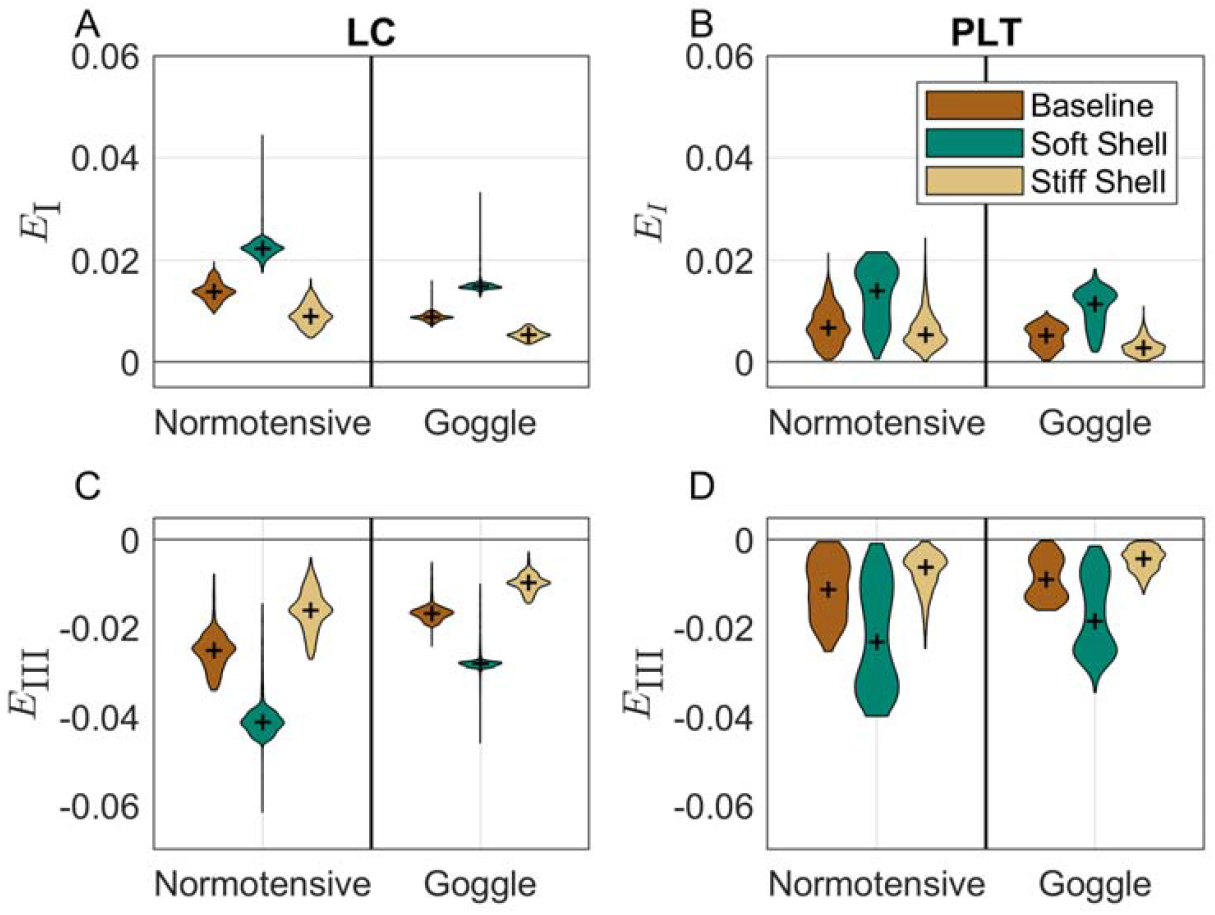
Sensitivity of strains in the LC (**A** and **C**) and PLT (**B** and **D**) to changing the corneoscleral shell stiffness, without fiber reinforcement in the ppSC. Contrary to the sensitivity to corneoscleral stiffness cases when considering a fiber-reinforced ppSC (Figure 4 **A, B, E**, and **F**), decreasing (increasing) the corneoscleral stiffness by a factor of two increased (decreased) LC and PLT strain magnitudes (*E_I_* and *E_III_*) by a factor of 1.6-2.1 (0.6-0.8).

**Figure S4:**
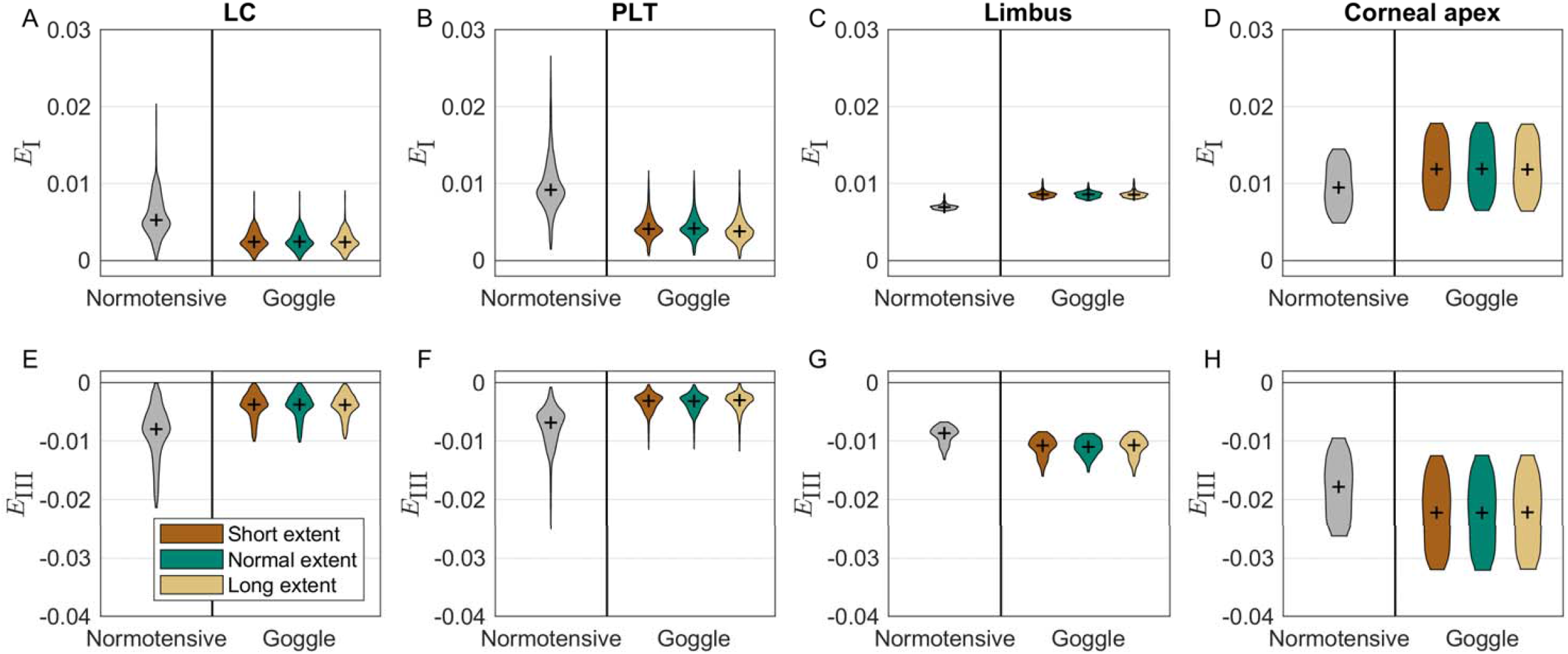
Effects of changing the spatial distribution of NPP on tissue strains, specifically **E_I_**(**A-D**), and (**E-H**) in the LC (**A** and **E**), the PLT (**B** and **F**), the limbus (**C** and **G**), and at the corneal apex (**D** and **H**). In the first variation, NPP was spatially uniform and only affected the anterior aspect of the eye from the corneal apex to ∼2.5 mm posterior to the limbus (“NPP Short extent”). In the second case (“NPP Normal extent”), NPP was applied from the corneal apex to ∼2.5 mm posterior to the limbus, then decayed linearly to the anterior margin of the ppSC. In the third (“NPP Long extent”), NPP was specified to be spatially uniform from the corneal apex to the anterior margin of the ppSC, i.e. without the linear decrease of the NPP Normal extent case. Changing the NPP distributions had virtually no effect on or Data is shown using violin plots and the median is marked with ‘+’.

